# The intensities of canonical senescence biomarkers integrate the duration of cell-cycle withdrawal

**DOI:** 10.1101/2023.03.18.533242

**Authors:** Humza M. Ashraf, Brianna Fernandez, Sabrina L. Spencer

## Abstract

Senescence, a state of permanent cell-cycle withdrawal, is difficult to distinguish from quiescence, a transient state of cell-cycle withdrawal. This difficulty arises because quiescent and senescent cells are defined by overlapping biomarkers, raising the question of whether quiescence and senescence are truly distinct states. To address this, we used single-cell time-lapse imaging to distinguish slow-cycling quiescent cells from *bona fide* senescent cells after chemotherapy treatment, followed immediately by staining for various senescence biomarkers. We found that the staining intensity of multiple senescence biomarkers is graded rather than binary and primarily reflects the duration of cell-cycle withdrawal, rather than senescence per se. Together, our data suggest that quiescence and senescence are not distinct cellular states but rather fall on a continuum of cell-cycle withdrawal, where the intensities of canonical senescence biomarkers reflect the likelihood of cell-cycle re-entry.

## Introduction

Senescence is a state of permanent cell-cycle withdrawal associated with aging and DNA damage. Extended durations of recovery from DNA damaging chemotherapy treatments lead to cell-cycle re-entry and population regrowth (1,2), but it is unknown whether this regrowth phenotype is caused by cells that re-enter the cell cycle from a reversible state of arrest called quiescence or whether it is the result of a proliferative subpopulation that outcompetes senescent cells over time (Fig. 1A). It is challenging to study transient vs. permanent cell-cycle withdrawal since these fates are nearly indistinguishable from each other at a single point in time, making it unclear which cells will go on to cycle in the future vs. which cells will continue to remain arrested (3). These limitations have led to speculation that some cells can escape from senescence to re-enter the cell cycle (2,4), but it has not been shown that these cells were truly senescent to begin with. As a result, there is a critical need to accurately detect senescent cells to clarify whether quiescence and senescence are binary, distinct cellular states or whether they exist on a gradient of cell-cycle withdrawal.

**Figure 1.**
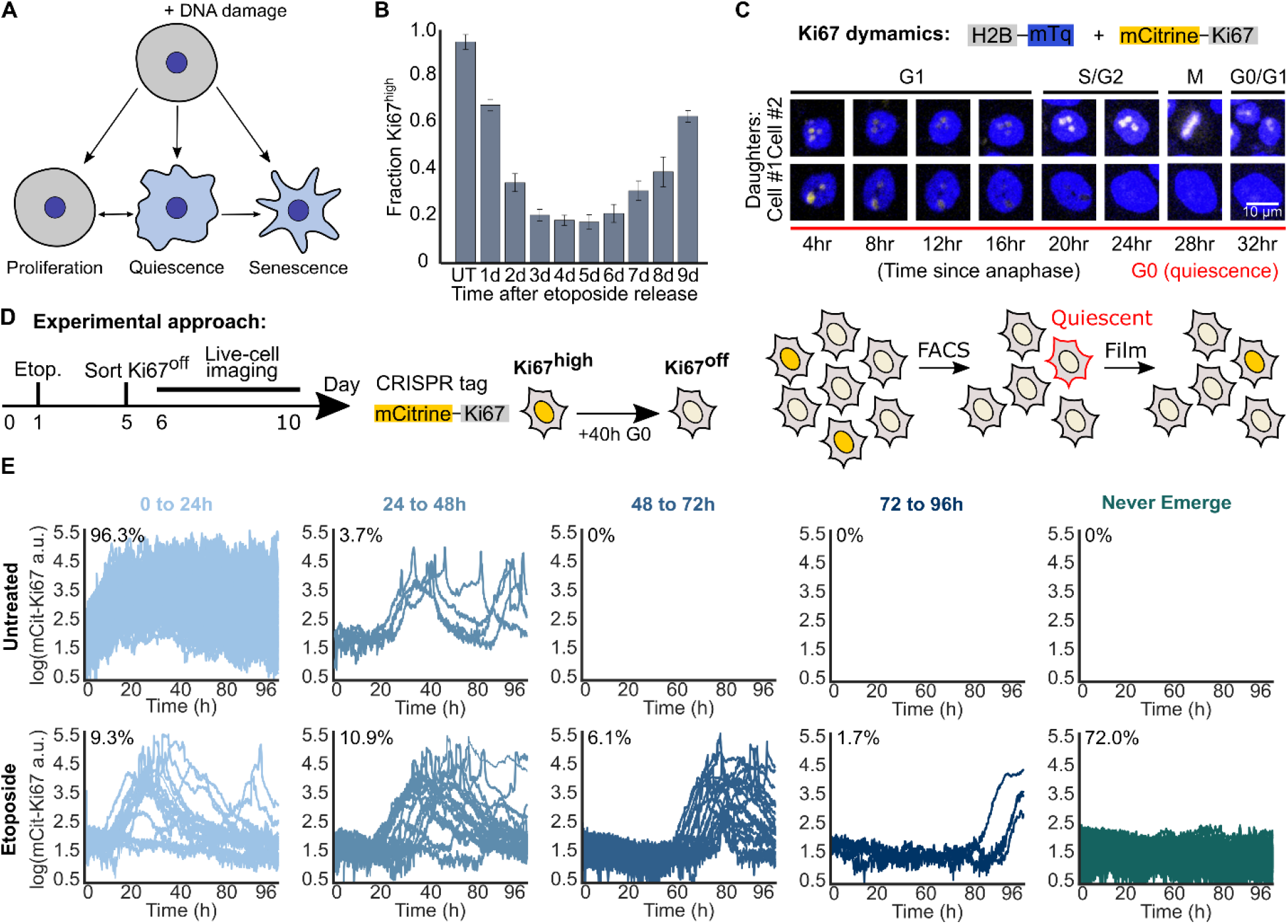
A subpopulation of cells exits quiescence to re-enter the cell cycle after chemotherapy treatment. (A) Multiple cell fates arise during recovery from acute DNA damaging agents. (B) MCF10A cells were treated with 10 μM etoposide for 24h before being washed and allowed to recover for 1 to 9d. Cells were fixed and stained for Ki67, and the fraction of Ki67^off^ cells was calculated for each condition. (C) Dynamics of mCitrine-Ki67 with respect to cell cycle phase in two daughter cells originating from the same mother. (D) Experimental schematic: MCF10A cells expressing endogenously tagged with mCitrine-Ki67 were plated on day 0 and treated on day 1 with 10 μM etoposide for 24h and washed. On day 5, Ki67^off^ cells were isolated by flow cytometry, plated, and allowed to grow for 24h before being imaged for 96h by time-lapse microscopy. (E) Single-cell traces are grouped based on their relative timing of cell-cycle re-entry from the Ki67^off^ state and the percent of cells in each group is indicated. 200 cell traces total are plotted in each row.

Since no single senescence marker is unique to senescence, multiplexing multiple markers in single cells has been suggested as a new goal to identify senescent cells more accurately (3,5). The gold-standard marker for detecting senescent cells is the senescence-associated-beta-galactosidase (SA-β-Gal) stain (3,6); however, the β-galactosidase gene is dispensable for the induction and maintenance of senescence (7), raising questions about a causal relationship between SA-β-Gal positivity and irreversible cell-cycle withdrawal. Furthermore, there is no robust method for quantifying SA-β-Gal. Most studies simply binarize the colorimetric stain by manually labelling cells either blue (senescent) or not blue (not senescent). Due to these limitations, studies often measure other senescence markers in parallel experiments. These include the lack of cell-cycle markers (e.g. Ki67 and phospho-Rb), expression of Cyclin-Dependent Kinase (CDK) inhibitors (e.g. p21 and p16), DNA damage (e.g. 53BP1 or γH2AX), presence of the senescence-associated-secretory phenotype (SASP, with IL6 as one of the most common factors), and increased cell size (3,5). However, no study has systematically tested these markers against a ground-truth readout of senescence to quantify their predictive power for identifying senescent cells.

Here, we developed a ground-truth readout of senescence using long-term single-cell time-lapse imaging of cell-cycle markers. Using our approach, we classified *bona fide* senescent cells as those that never entered the cell cycle during the course of a four-day movie and distinguished them slow-cycling quiescent cells. We mapped these cell-cycle behaviors to post hoc SA-β-Gal staining by developing a novel method for quantifying and multiplexing the stain with other senescence biomarkers. We found that the relative blueness of the SA-β-Gal stain reflects increased lysosomal content and scales with increasing durations of cell-cycle withdrawal, rather than senescence per se. Furthermore, the intensity of several other markers also scales with the duration of cell-cycle withdrawal, including LAMP1, 53BP1, and cytoplasmic and nuclear area. Together, our data suggest that that quiescence and senescence are not distinct cellular states but rather fall on a continuum of cell-cycle withdrawal, where the likelihood of cell-cycle re-entry is strongly correlated with the relative intensities of canonical senescence biomarkers.

## Results

### A subset of cells re-enters the cell cycle from quiescence and contributes to population regrowth following chemotherapy treatment

To measure the heterogeneity in cell-cycle fates following senescence induction, we treated MCF10A non-transformed mammary epithelial cells with etoposide, a commonly used anti-cancer chemotherapeutic agent that inhibits topoisomerases to induce DNA damage and senescence (8). MCF10A cells were released for 1-9d from a 24h treatment of 10 μM etoposide, and cells were fixed and stained for the proliferation marker Ki67 (**Fig. 1B and Fig. 1C**). Over the duration of recovery from drug, the majority of cells initially withdrew from the cell cycle, but the population began to rebound starting at day 6. This proliferation rebound was confirmed in MCF10A cells with an alternative proliferation marker (Rb phosphorylation, a cell-cycle marker that turns on when cells commit to the cell cycle and turns off when cells exit the cell cycle) (9), as well as in RPE-hTERT, MCF7, and WI38-hTERT cells (**Fig. S1A-C**). We observed further decreases in the proportion of proliferating cells at increasing doses of etoposide, but there was no concentration of drug (up to 50 μM) that eliminated all cycling cells to yield a pure senescent population (**Fig. S1C**).

The proliferating cells rapidly overtake the non-cycling cells by day 9 after chemotherapy treatment, but it remains unclear what proportion of the non-cycling cells at a snapshot in time are quiescent rather than senescent. To address this question, we used MCF10A cells in which Ki67 was tagged at the endogenous locus with mCitrine (10) to isolate non-cycling Ki67^off^ cells by flow cytometry 5d after treatment with etoposide, as this was the time point with the fewest cycling cells (**Fig. 1D and Fig. 1C**). The levels of Ki67 protein decays with second order kinetics upon cell-cycle exit, hitting the floor of detection after 40h (10). Thus, Ki67^off^ cells at the time of sorting have been out of the cell-cycle (quiescent or senescent) for at least 40h. Immediately after sorting, we replated the Ki67^off^ cells and began filming them the following day for an additional four days. 100% of the untreated, rare spontaneously quiescent Ki67^off^ cells re-entered the cell cycle within the first two days of filming, consistent with their quiescent status at the time of sorting. Surprisingly, despite the strong senescence-inducing conditions, 28% of etoposide-released cells resumed proliferation at some point during live-cell imaging (**Fig. 1E**). Similar results were obtained when this experiment was repeated with 10 Gy ionizing radiation (**Fig. S1D**). Thus, a significant fraction of non-cycling cells 5d after chemotherapy or ionizing-radiation treatment are actually quiescent and not senescent, since they are fated to re-enter the cell cycle in the future.

### SA-β-Gal staining overlaps between quiescent and senescent cell fates

Having developed a flow cytometry and time-lapse approach to identify *bona fide* senescent cells, we sought to clarify the relationship between cell-cycle withdrawal and SA-β-Gal staining, the gold-standard marker of senescence. To address the critical need in the senescence field for quantification of the SA-β-Gal stain, we adapted an existing method (11) to develop an automated, high-throughput strategy for measuring SA-β-Gal in thousands of single-cells (see Methods). We used the red component of the RGB image of the stain to compute a single value from the distribution of pixels within the cytoplasm of each segmented cell. We chose the red channel because the SA-β-Gal stain is primarily composed of blue and green pigments that preferentially absorb red light. Thus, the SA-β-Gal stain can be most easily quantified as the absence of red signal within every cell, and this channel has the largest dynamic range relative to background (**Fig. 2A**). We used the value at the 5^th^ percentile of red pixels within the cytoplasm of each cell as the SA-β-Gal score (**Fig. S2A-B**), since this method visually matched the relative blueness of cells (**Fig. S3A**) and recapitulated the gradient of staining in single cells induced to senescence (**Fig. 2A-B**). Despite the fact that SA-β-Gal is almost always used as binary marker of senescence, we found that the distribution of the SA-β-Gal signal is graded rather than binary, with no clear cut-off for designating a cell as senescent (**Fig. 2B**).

**Figure 2.**
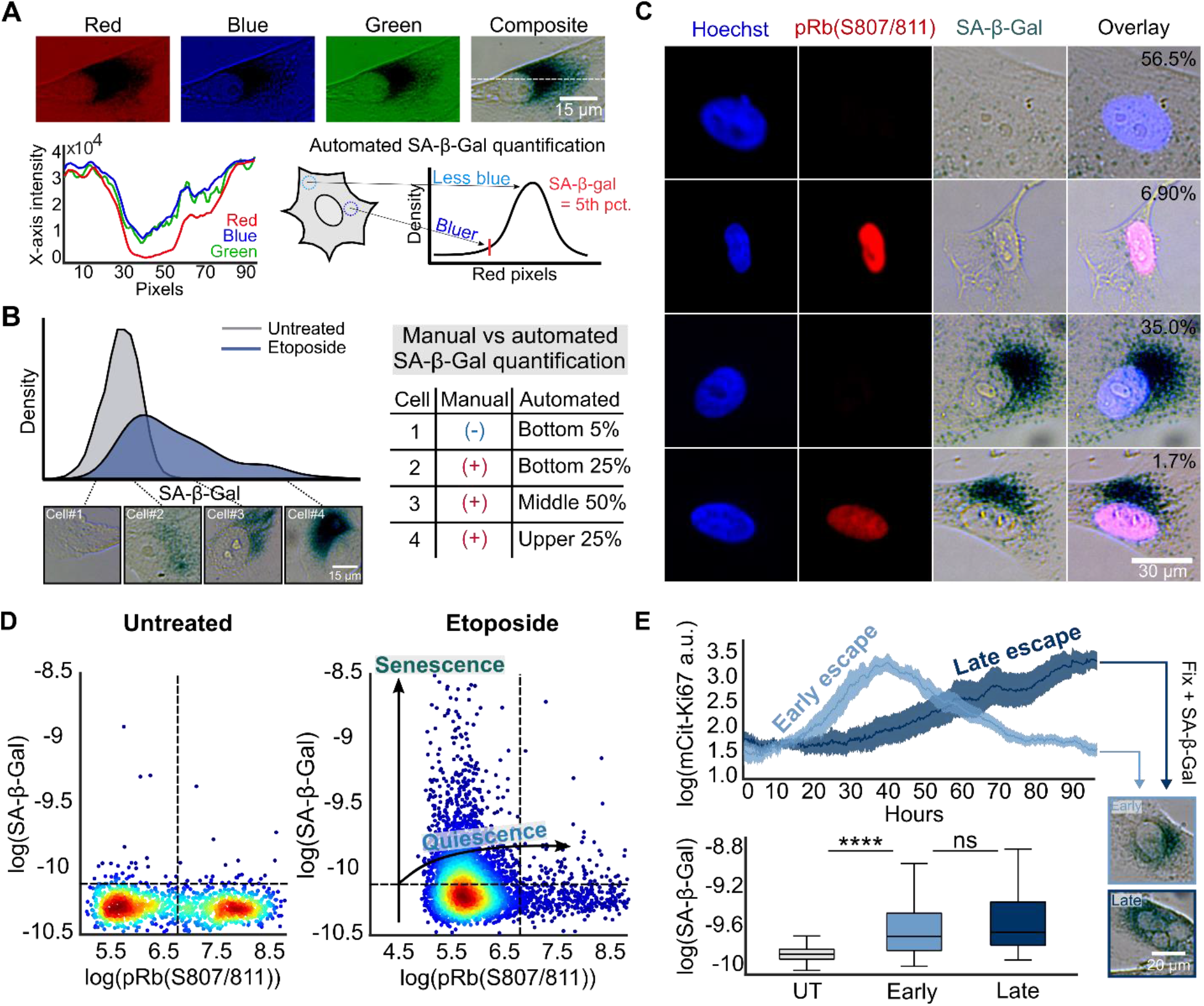
Quantifying SA-β-Gal in single cells shows significant overlap in staining between quiescent and senescent cells. (A) A representative single MCF10A cell stained for SA-β-Gal and imaged in pseudo-color-brightfield at 6d after release from 10 μM etoposide for 24h. An intensity profile (dotted line) was taken from each channel of the RGB stack. We define the SA-β-Gal signal as the 5^th^ percentile of the red pixels in the cytoplasm of each cell. (B) Distribution of SA-β-Gal signal after etoposide release with four representative single cells at increasing intensities of staining. (C) Heterogeneity in co-staining of SA-β-Gal and Rb phosphorylation by immunofluorescence in MCF10A cells released for 6d from a 24h treatment with 10 μM etoposide. Percentages reflect the fraction of cells with each behavior. (D) Scatter plots of SA-β-Gal versus phospho-Rb from the same cells in Fig. 2C. The arrows indicate the hypothesized quiescence versus senescence cell fate trajectories based on the relative level of SA-β-Gal staining. (E) The same data in Fig. 1D for etoposide-released cells that entered the cell cycle during live-cell imaging. Early escaping cells were those that were Ki67^off^ on the final frame of the movie while late escaping cells were those that were Ki67^high^ on the final frame of the movie. Single-cell traces were linked back to their relative SA-β-Gal levels after being fixed and stained for the marker at the end of imaging.

Next, we co-stained cells with SA-β-Gal and an antibody against Rb phosphorylation, a marker of cell cycle commitment (9), and discovered a surprisingly heterogeneous mixture of behaviors. Because the SA-β-Gal signal is graded and not binary (**Fig. 2B**), we initially used a cutoff at the 95^th^ percentile of untreated cells to designate a cell as SA-β-Gal-positive (hereafter SA-β-Gal^pos^). Although most SA-β-Gal^pos^ cells were phospho-Rb^low^, consistent with what would be expected for senescent cells, we also identified SA-β-Gal^neg^/phospho-Rb^high^ cycling cells, SA-β-Gal^neg^/phospho-Rb^low^ presumably quiescent cells, and a small fraction (1.7%) of unexpected SA-β-Gal^pos^/phospho-Rb^high^ cells. This latter population calls into question the reliability of SA-β-Gal as a senescence marker, since no truly senescent cell should ever be in the cell-cycle (**Fig. 2C**). However, comparing the relative intensities of SA-β-Gal staining following etoposide release revealed that the bluest cells in the population were significantly more likely to be phospho-Rb^low^ compared to less-blue cells, which were associated with more variability in phospho-Rb status (**Fig. 2D**). This suggests that the confidence in classifying cells as senescent increases as a function of the intensity of SA-β-Gal staining, with intermediate levels of SA-β-Gal staining encompassing both reversibly and irreversibly arrested cells.

To determine the origin of the SA-β-Gal^pos^/phospho-Rb^high^ cells, we returned to our data set from Fig. 1D-E where the cells were also stained cells for SA-β-Gal at the end of the movie. Because SA-β-Gal^pos^/phospho-Rb^high^ cells tended to have intermediate levels of SA-β-Gal staining, we hypothesized that this subpopulation might represent slow-cycling cells that we showed to be easily misclassified as senescent (**Fig. 1D**). To test this, we split the slow cycling subpopulation that re-entered the cell cycle during imaging into two categories: early vs. late escapers, based on whether the cells were Ki67^off^ or Ki67^high^ on the final frame of the movie (**Fig. 2E, top**). As expected, we observed slow cycling cells that happened to be in the cell cycle at the final frame of the movie to have significantly higher levels of SA-β-Gal staining compared to untreated control cells, explaining the origin of the SA-β-Gal^pos^/phospho-Rb^high^ subpopulation (**Fig. 2E, bottom**). However, we detected no significant difference in the relative levels of blueness between the early and late escaping subpopulations, suggesting that past proliferative history rather than current cell-cycle status determines the eventual levels of SA-β-Gal staining (**Fig. 2E, bottom**). Together, these data support two major conclusions: 1) cell cycle re-entry does not immediately extinguish SA-β-Gal staining, explaining the SA-β-Gal^pos^/phospho-Rb^high^ subpopulation, and 2) SA-β-Gal positivity marks both reversibly and irreversibly arrested cell fates, rather than exclusively cellular senescence.

### SA-β-Gal intensity scales with increased durations of cell-cycle withdrawal

To determine whether the overlap in SA-β-Gal staining between quiescent and senescent cells is due to the fact that its signal rises as a function of the duration of cell-cycle withdrawal, we filmed cells expressing a live-cell sensor for CDK2 activity (12) for days 5-9 following etoposide release (**Fig. 3A**). CDK2 activity begins to rise when cells commit to the cell cycle and increases steadily thereafter until mitosis, whereas cells turn off their CDK2 activity and enter a CDK2^low^ state when they exit the cell cycle. At the end of the time-lapse imaging on day 9, we fixed and stained the cells for SA-β-Gal and mapped each cell’s stain to its cell-cycle history over the previous four days (**Fig. 3B**). Binning cells into the top, middle, and bottom 10% of SA-β-Gal signal revealed that the intensity of staining was proportional to the total duration of time that cells spent out of the cell cycle in a CDK2^low^ state. This shows that SA-β-Gal staining scales with increasing durations of cell-cycle withdrawal (**Fig. 3C, left**).

**Figure 3.**
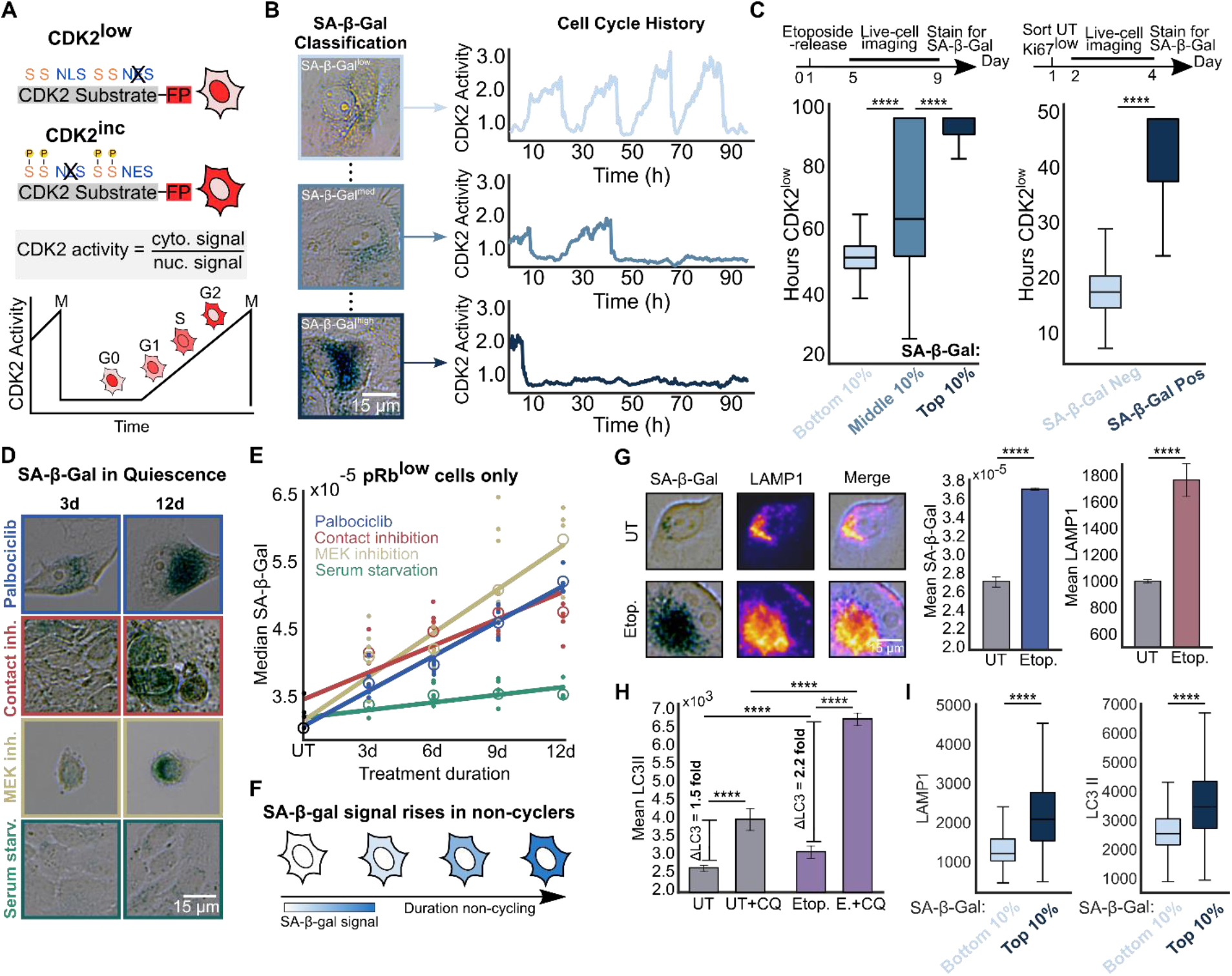
SA-β-Gal staining marks long durations of cell cycle exit and is correlated with increased lysosomal content and autophagic flux. (A) Schematic of the CDK2 activity sensor. The sensor localizes to the nucleus when unphosphorylated; progressive phosphorylation by CDK2 leads to translocation of the sensor to the cytoplasm. NLS, nuclear localization signal; NES, nuclear export signal; S, CDK consensus phosphorylation sites on serine. (B) MCF10A cells expressing the CDK2 activity sensor were treated with 10 μM etoposide for 24h, washed, and subjected to time-lapse microscopy 5d later for 96h (from 5d-9d). The cells were fixed and stained for SA-β-Gal after the last frame was taken; (C: left) single-cell traces were clustered based on the top, middle, and bottom 10% of SA-β-Gal signal and the total hours CDK2^low^ (below a cutoff of 0.8) was plotted for each bin. (C: right) Untreated MCF10A cells were sorted by flow cytometry for the bottom 1% of mCitrine-Ki67 signal, plated and allowed to grow for 48h, filmed for 48h to monitor CDK2 activity, fixed and stained for SA-β-Gal, and were manually classified as SA-β-Gal positive versus negative. (D-E) MCF10a cells were pushed into quiescence by contact inhibition, serum starvation, 3 μM palbociclib treatment, or 100 nM trametinib treatment for 3-12d and fixed and stained for SA-β-Gal and phospho-Rb. Best fit lines were computed for each condition from the average of 6 technical replicates. (F) Model for SA-β-Gal accumulation as a function of cell-cycle exit time. (G) MCF10A cells were treated with 10 μM etoposide for 24h, washed, fixed after 3d, and stained for SA-β-Gal and LAMP1. (H-I) Same experimental scheme as described in Fig. 3G. Cells were fixed and stained for SA-β-Gal, LC3II, and LAMP1 after a 3 h treatment with 50 μM chloroquine.

To test whether SA-β-Gal also has the resolution to identify cells withdrawn from the cell cycle due to low-grade endogenous stress (12), we quantified the intensity of SA-β-Gal staining in spontaneously quiescent cells in an unperturbed population. We sorted the bottom 1% of mCitrine-Ki67 signal by flow cytometry to enrich for the intrinsically slow-cycling cells in the population (**Fig. 3C right**). The cells were re-plated after sorting and their CDK2 activities were filmed over the subsequent two days. Cells designated SA-β-Gal^pos^ spent significantly more hours in the CDK2^low^ state compared to SA-β-Gal^neg^ cells, supporting the notion that SA-β-Gal is a general readout of increased durations of cell-cycle withdrawal regardless of the stressor.

To extend these findings to multiple types of cell-cycle withdrawal, we measured the median SA-β-Gal signal in non-cycling cells after forcing cells into quiescence for 3, 6, 9, and 12d using four well-established methods: contact inhibition, serum starvation, CDK4/6 inhibition (Palbociclib), and Mek inhibition (Trametinib) (**Fig. 3D**). Unlike etoposide, these treatments induce a transient cell-cycle exit that reverses after washout (13), so increases in SA-β-Gal staining within the non-cycling cells stems from quiescent rather than senescent cells. Across all four treatment conditions, the SA-β-Gal signal increased as a function of treatment time (**Fig. 3E and Fig. S4),** supporting the notion that SA-β-Gal is a graded marker whose signal intensity integrates the duration of time spent out of the cell cycle (**Fig. 3F**). We conclude that heterogeneity in SA-β-Gal staining is reflective of biological heterogeneity, where cells that cycle less often under stress accumulate more SA-β-Gal staining over time.

### Increased SA-β-Gal staining reflects increased lysosomal content and autophagy

Why is SA-β-Gal staining is so closely coupled with cell-cycle status when the enzyme itself is dispensable for the induction and maintenance of senescence (7,14)? Since the β-galactosidase enzyme is localized to the lysosomes, we reasoned that increased SA-β-Gal staining could simply be a readout of increased lysosomal content. To test this, we multiplexed measurements of SA-β-Gal and LAMP1, a membrane-embedded lysosomal protein, and found that they not only co-localized but the levels of both also simultaneously increased following acute chemotherapeutic stress (**Fig. 3G**). This suggests that increased lysosome biogenesis following cell stress leads to increased activity of β-galactosidase.

We next questioned whether the increased lysosome content following etoposide release was linked to changes in autophagy, since previous literature has suggested that senescent cells undergo increased autophagic flux to manage the accumulation of cellular damage (15). To investigate this idea, we compared the autophagic flux in untreated and etoposide-released cells by comparing the relative increase in LC3II protein levels following a 3h treatment of 50 μM chloroquine (CQ), a lysosomotropic agent that impairs autophagosome fusion (16). Etoposide-released cells experienced a 2.2-fold increase in average LC3II protein levels following CQ treatment compared to control cells, which had a 0.5-fold increase. This result suggests that autophagy is significantly upregulated in cells released from acute chemotherapeutic stress (**Fig. 3H**).

To test whether increased SA-β-Gal staining is correlated with both increased lysosomal content and/or autophagic flux, we co-stained cells for SA-β-Gal and either LAMP1 or LC3II following etoposide release. As expected, the levels of both proteins were more significantly upregulated in cells with the highest levels of SA-β-Gal staining compared to cells with the lowest (**Fig. 3I**). Thus, SA-β-Gal staining reflects increased lysosomal content, which reflects increased autophagic flux in cells induced to senescence.

### Canonical senescence biomarkers resolve cycling from non-cycling cells better than they resolve quiescent from senescent cells

Because SA-β-Gal staining scaled with increased durations of cell-cycle withdrawal, we next asked whether other markers of senescence follow the same trend. To test this, we simultaneously multiplexed SA-β-Gal, LAMP1, cytoplasmic area, and 53BP1, a protein that forms large nuclear bodies at sites of DNA damage (17), after 4d of live-cell imaging, beginning 5d after etoposide release. First, we classified cells as either proliferative, quiescent, or senescent based on the number of hours spent in the CDK2^low^ state (**Fig. 4A),** where senescent cells were defined as cells that were CDK2^low^ for the entire movie (see **Fig. S5** for classification). For all four markers, the intensity of staining was highest for senescent cells, intermediate for quiescent cells, and lowest for proliferative cells (**Fig. 4B**). Second, we grouped cells based on marker staining intensity into the top, middle, and bottom 10% and plotted the time the cells had spent in the CDK2^low^ state over the prior 4 days. For all four markers tested, the same graded trend was observed, where the intensity of staining of the marker was correlated with the duration of time withdrawn from the cell cycle. (**Fig. 4C**). These data suggest that the relative intensities of canonical senescence biomarkers encode in a snapshot information about the cell-cycle histories of single cells released from acute chemotherapeutic stress.

**Figure 4.**
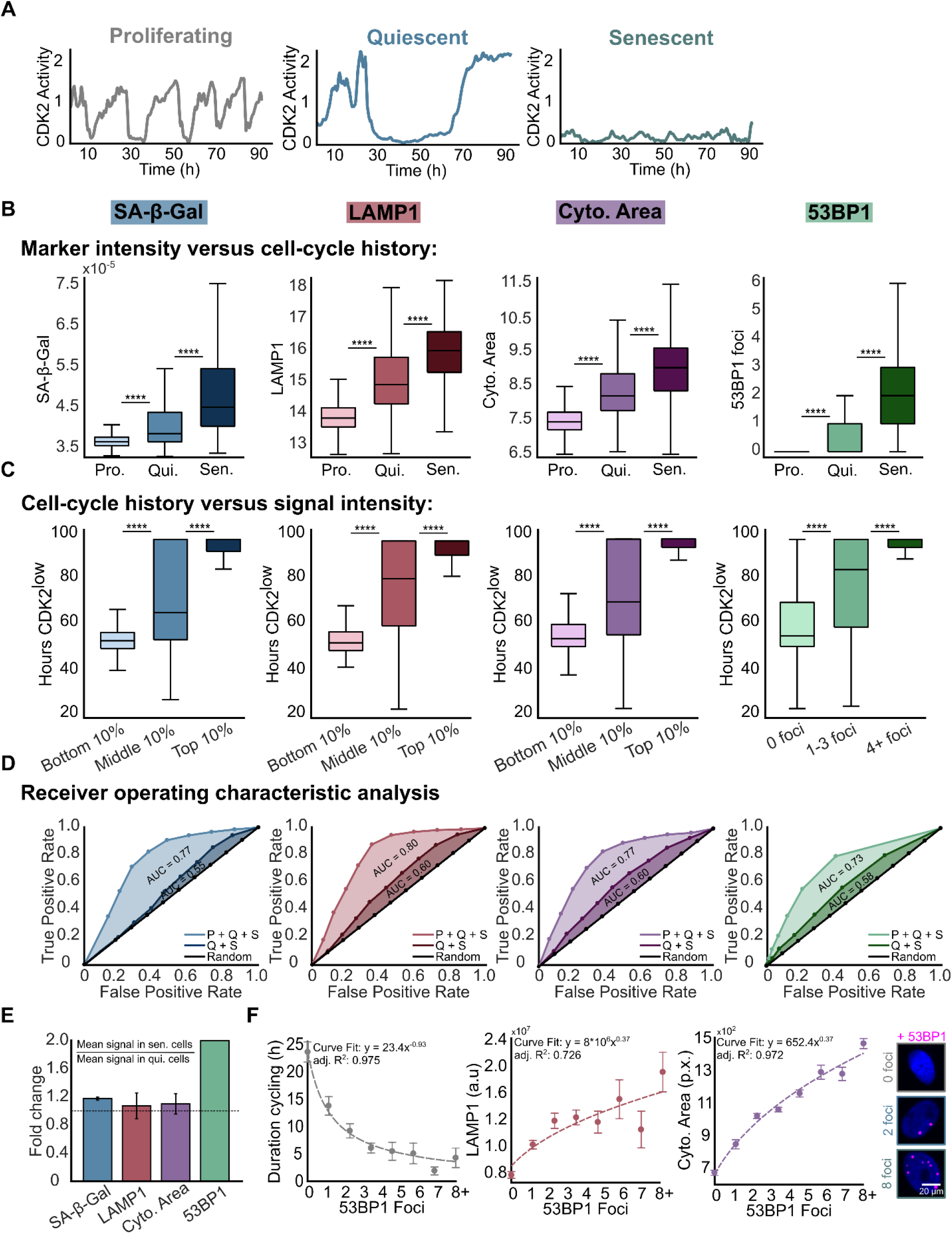
Senescence biomarkers generally resolve cycling from non-cycling cells better than quiescent from senescent cells. (A-F) MCF10A cells expressing the CDK2 activity sensor were treated with 10 μM etoposide for 24h, washed, and subjected to time-lapse microscopy 5d later for 96h (from 5d-9d). The cells were fixed and stained for SA-β-Gal, LAMP1, succinimidyl ester, and 53BP1 after the last frame was taken. (A-B) Cells were split into proliferative, quiescent, or senescent based on their duration spent CDK2^low^ during live cell imaging, and the intensity of each marker was measured for each cellular fate. (C) The duration cells spent in the CDK2^low^ state was plotted against the bottom, middle, and top 10% of signal for each marker. (D) ROC analysis for proliferative, quiescent, and senescent cells versus only quiescent and senescent cells. AUC indicates the area under the curve for each condition. (E) Fold changes of the mean intensity of senescence markers in senescent versus quiescent cells. (F) The duration spent CDK2^high^, LAMP1 intensity, or cytoplasmic area versus 53BP1 foci number from the same experiment as described in Fig. 4A-E.

To quantitatively compare the power of these senescence markers to accurately identify senescent cells, we generated receiver operating characteristic (ROC) curves for each of the markers. The ROC curve compares the true positive rate versus the false positive rate at increasing thresholds of detection, where high thresholds maximize true positives and low thresholds minimize false negatives. In this case, we classified cells as true positives if they remained CDK2^low^ throughout the duration of the movie, which we defined as the ground-truth senescent subpopulation. For each marker, we computed two ROC curves: the first was for all cells in the population while the second was for only quiescent and senescent cells. This analysis allowed us to compare the relative resolving power for each senescence biomarker to differentiate 1) cycling from non-cycling cells and 2) non-cycling quiescent from non-cycling senescent cells. Unsurprisingly, removing the proliferative subpopulation from the analysis significantly reduced the ability to accurately classify cells as senescent (**Fig 4D**). Thus, non-cycling (quiescent and senescent) cells are more easily resolved from cycling cells than quiescent and senescent cells can be resolved from each other.

### The extent of DNA damage after etoposide release dictates the probability of cell-cycle re-entry

To determine which single senescence marker had the best ability to separate quiescent from senescent cells at a snapshot, we computed the fold change in signal for SA-β-Gal, LAMP1, cytoplasmic area, and 53BP1 in quiescent versus senescent cells (**Fig. 4E**) using the dataset from Fig 4A-D. This analysis revealed that 53BP1 was the most enriched in senescent cells compared to quiescent cells, while every other marker overlapped significantly between these two states (**Fig. 4E**). We therefore measured whether increasing numbers 53BP1 nuclear bodies directly correlated with increases in other markers of senescence including reduced proliferation, LAMP1 staining, and cytoplasmic area (**Fig. 4F**). For each marker, we observed a stepwise change in mean intensity as a function of 53BP1 (**Fig. 4F**), suggesting that the extent of DNA damage (and not simply the binary presence or absence of DNA damage) is a critical determinant of irreversible cell cycle withdrawal. We speculate that this is a consequence of increased DNA damage causing a concurrent increase in tumor suppressor genes that block cell-cycle entry, such as p21 (1).

### The largest cells in the population go on to accumulate features of truly senescent cells

Increased cell size is one of the first identified and most universally upregulated markers of cellular senescence (18). Recent work has shown that the molecular features associated with senescent cells are directly caused by aberrant increases in cell size (19). To test to what extent cell size could separate quiescent from senescent cells prior to cell-cycle re-entry following etoposide release, we returned to our dataset from Fig. 1D where untreated and etoposide-released mCitrine-Ki67 MCF10A cells were sorted to be Ki67^off^ and replated for timelapse imaging the following day. Cells were classified as proliferating (untreated yet Ki67^off^ at the time of sorting), quiescent (etoposide-released and re-entered the cell cycle before the movie ended), or senescent (etoposide-released and remained non-cycling Ki67^off^ throughout the movie), and their mean nuclear areas were compared in the first 10h of filming when every cell was still Ki67^off^ (**Fig 5A**). Senescent cells were significantly larger than quiescent cells in this 10h window preceding escape from the Ki67^off^ state (**Fig 5A, right**), suggesting that increased cell size is correlated with a reduced likelihood for cell-cycle re-entry.

**Figure 5.**
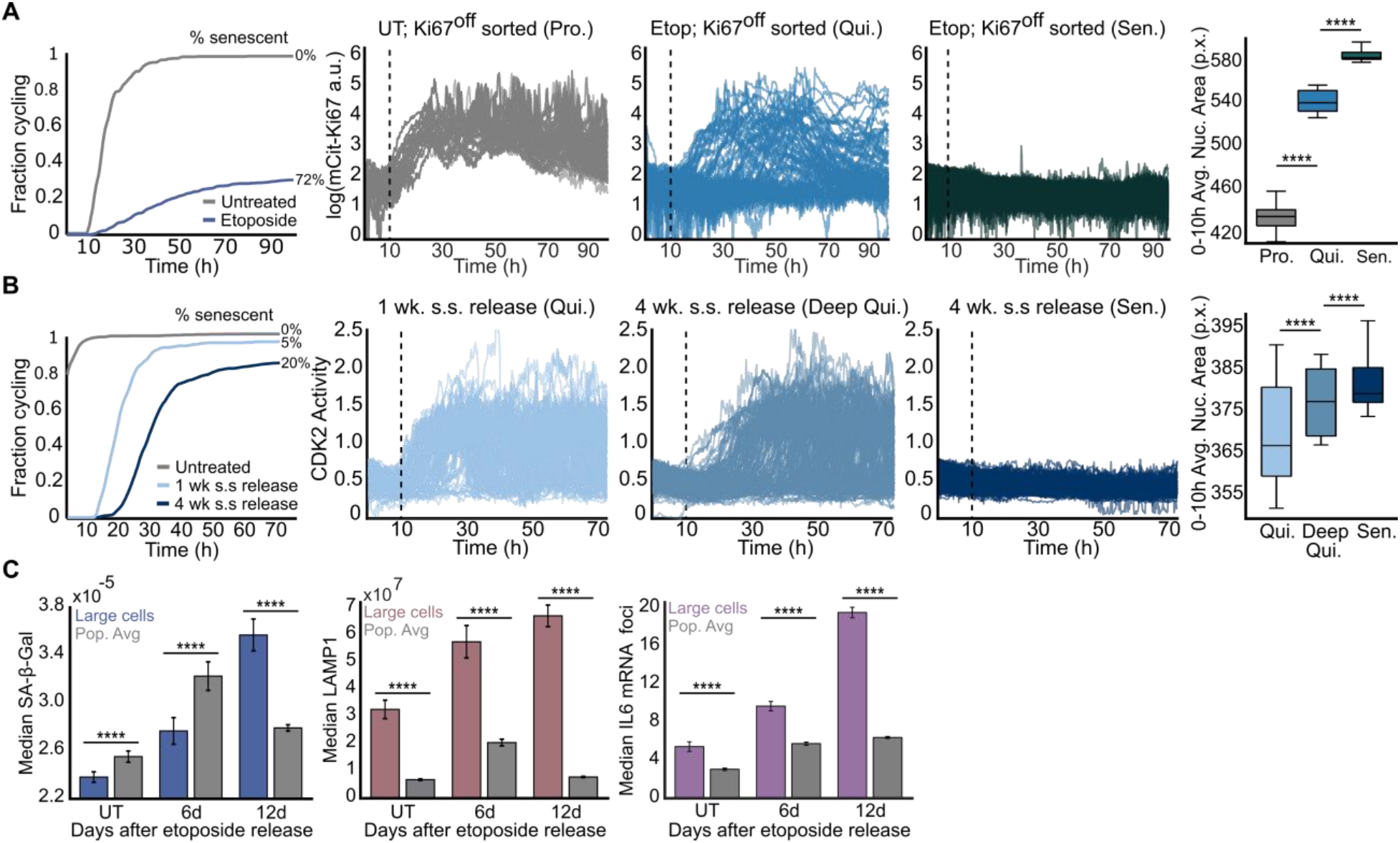
The largest cells in the population go on to accumulate features of truly senescent cells. (A) The data as in Fig. 1D. Cumulative distribution function (CDF) curves represent the fraction of cells that re-enter the cell cycle during the movie; the remainder of cells are defined as senescent (72%). Cells were clustered based on their relative mCitrine-Ki67 signals: proliferating (untreated; gray); quiescent (etoposide released; blue); senescent (etoposide released; green). The mean nuclear area within the first 10h of filming was plotted for each category. (B) Live-cell imaging of CDK2 activity after serum re-addition for cells serum starved for 1 or 4 weeks. CDF curves represent the fraction of cells that re-enter the cell cycle during the movie; the remainder of cells are defined as senescent (5% after 1 week of serum starvation and release, 20% after 4 weeks). The 4-week condition was split into cells that re-entered the cell cycle during the movie (deep quiescence) versus those that did not (senescent). The mean nuclear area within the first 10h of filming was plotted for each category. (C) SA-β-Gal, LAMP1, and IL6 mRNA signal was measured over the course of a 12d recovery after release from a 24h pulse of 10 μM etoposide treatment in all cells versus cells in the top 10% of cytoplasmic areas.

Due to the phenotypic similarities between quiescent and senescent cells, we next asked whether increased cell size could mark the transition from quiescence to senescence as a function of cell-cycle withdrawal time. Because increased durations of time spent in quiescence impairs the ability of cells to re-commit to the cell cycle (20), we serum starved MCF10A cells for one and four weeks and filmed the timing of cell cycle re-entry after serum re-stimulation. Four-week starved cells not only took longer to re-enter the cell cycle compared to one-week starved cells but also had a significant fraction (20%) that became senescent **(Fig. 5B)**. We plotted the mean nuclear area of each cell subpopulation in the first 10h of filming as in Fig. 5A and found that larger cell sizes correlate with delayed cell-cycle re-entry, albeit with significant overlap in the size distributions for each cellular fate (**Fig. 5B, right**). These data support the notion that reversible and irreversible cell-cycle arrest exist on a continuum.

Finally, because the largest cells in the population at a snapshot are more likely to remain arrested into the future, we questioned whether increased cell size was linked to a higher intensity of staining of canonical senescence markers over time. To test this, we measured SA-β-Gal and LAMP1 at various timepoints following a 24h treatment of 10 μM etoposide and plotted the median signal for all cells as well as for cells falling within the top quartile of cytoplasmic area (**Fig. 5C, left and Fig. 5C, middle**). As expected, the SA-β-Gal and LAMP1 within the entire population rose and fell along the duration of recovery, as proliferating cells eventually outgrew non-cycling cells. However, within the subset of larger cells, SA-β-Gal and LAMP1 continuously increased over time. (**Fig. 5C, left and Fig. 5C, middle**). Furthermore, we performed the same analysis on cells stained for IL6, the most commonly upregulated SASP factor (21,22), by mRNA fluorescence in situ hybridization (FISH), and observed a similar trend (**Fig. 5C, right**). Thus, the largest cells following etoposide release accumulate a canonical senescent phenotype overtime, while the remainder of the population re-enters the cell cycle and contributes to population regrowth.

## Discussion

Measuring senescence with either a single marker or at a single point in time can lead to incorrect conclusions about the biology or dynamics of senescent cells. To address these gaps, we used time-lapse imaging to differentiate quiescent from senescent cells following acute chemotherapeutic stress, and we developed novel methods for multiplexing and quantifying senescence biomarkers in single cells. These data revealed that increasing durations of cell-cycle withdrawal correlate with increasing SA-β-Gal, LAMP1, cytoplasmic area, and 53BP1 levels and that multiplexing senescence biomarkers in single cells increased the resolution for detecting senescent cells over time.

Previous studies have reported that in certain contexts cells can escape from senescence to resume proliferation in the future (2,23); however, our time-resolved analysis of single cells induced to senescence suggests that this regrowth phenotype stems from cells that were never truly senescent. Instead, quiescent cells that retain their capacity to proliferate can outcompete true senescent cells over time in a heterogenous population. These cells may be confused as senescent since they can also stain positive for canonical senescence biomarkers such as SA-β-Gal, which we found to mark long durations of cell-cycle withdrawal rather than senescence *per se*. Furthermore, increased SA-β-Gal staining was associated with increased autophagic flux (**Fig. 3H-I**), a phenotype that was also observed in deeply quiescent cells, albeit at reduced efficiency (20). This is consistent with the fact that increased autophagy has been shown to be an activator of cellular senescence (15), suggesting that increased autophagy may be a general feature of stress-induced cell-cycle exit and may control the probability of cell cycle re-entry in the future.

When we measured the intensities of multiple senescence biomarkers following etoposide release, we found that SA-β-Gal staining was not the only marker that was linked to a decreased likelihood for proliferation. Indeed, increased cell size, LAMP1 staining, and 53BP1 foci number were also strongly correlated with a reduction in the proliferative fate of single cells (**Fig 4F)**. Among these markers, we were able to measure nuclear area prior to cell-cycle re-entry following etoposide-release or serum starvation-release via live-cell imaging and found that nuclear area correlates with future cell cycle commitment outcomes. This suggests that cell size is not only associated with whether cells will re-enter the cell cycle in the future but also encodes information about the duration of cell-cycle withdrawal in the past. These data are consistent with the emerging concept that increased cell size is causal for the induction of cellular senescence (19,24). Several classes of canonical senescence biomarkers, such as increased lysosomal content, superscale with cell size (19), explaining the strong correlation between cell-cycle exit and the intensities of senescence biomarker staining in the largest cells over time (**Fig. 5C**). Thus, we propose that the pathway to senescence following acute chemotherapeutic stress is initially caused by irreparable levels of DNA damage and then maintained by aberrant increases in cell size. Since quiescent cells have less damage compared to senescent cells (**Fig. 4E**), they re-enter the cell cycle at a high enough frequency that they fail to grow as large as senescent cells.

While our analysis suggests that cell-cycle withdrawal is on a continuum where the likelihood of cell-cycle re-entry is related to the relative levels of senescence markers (**Fig. 6**), there may be some unique molecular features associated with quiescent vs. senescent cells. Future studies with the ability to classify cells as reversibly vs. irreversibly arrested at a snapshot in time will be critical for establishing more universal definitions for these phenotypes. This would help clarify the functional importance of the quiescent subpopulation that emerges alongside senescent cells after chemotherapy treatment, since our study calls into question the link between irreversible arrest and expression of senescence markers.

**Figure 6.**
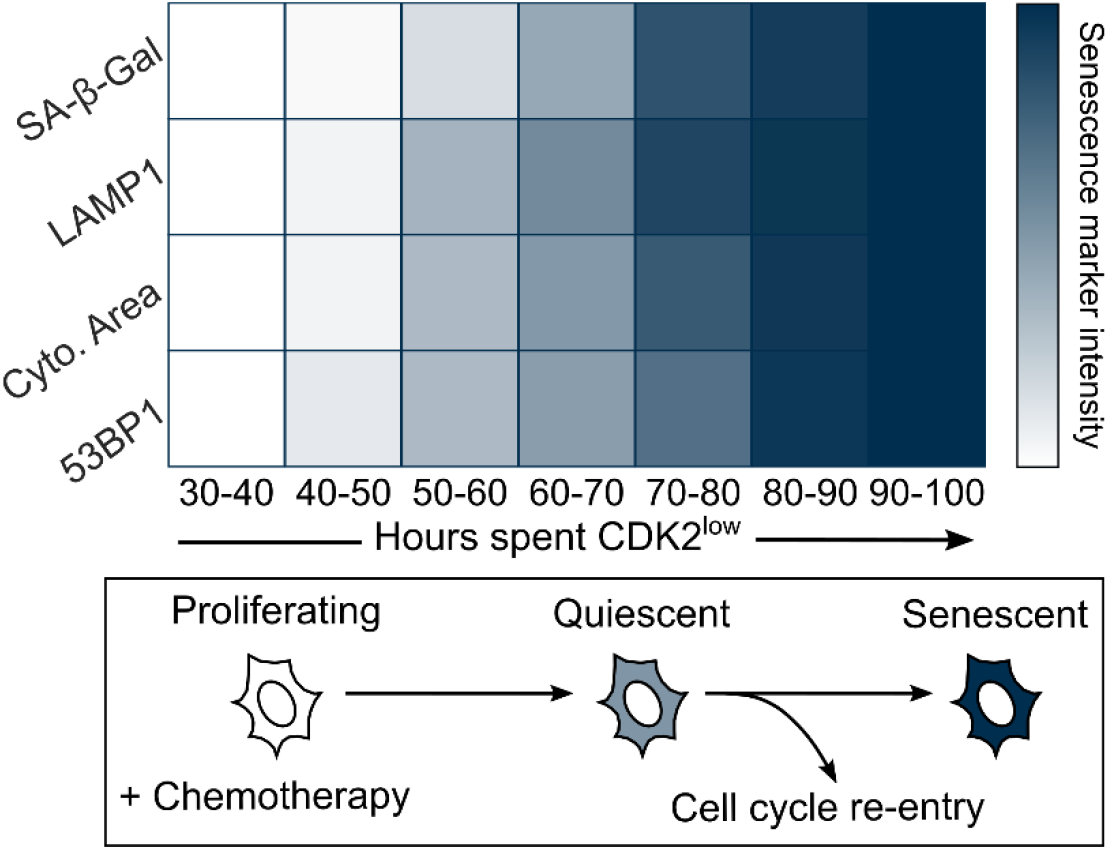
Summary of major findings. Cells were grouped based on the amount of time spent in the CDK2^low^ state from day 5-9 after etoposide release. The marker intensities in each group were averaged, scaled between 0 and 1, and each box was colored accordingly. This reveals a gradual, monotonic accumulation of each marker signal as a function of the duration of cell-cycle withdrawal. All data are from Fig. 4.

## Methods

### Antibodies and reagents

Anti-Ki67 (ab15580) and LC3 II (ab192890) were purchased from Abcam and used at 1:2000 and 1:1000 dilutions. pRb (S807/811) D20B12 XP (8516) and LAMP1 D2D11 XP (9091) were purchased from CST and used at 1:500 and 1:1000 dilutions. Anti-human 53BP1 (612523) was purchased from BD and was used at a 1:1000 dilution. LAMP1 (sc-20011) was purchased from Santa Cruz Biotech and used at a 1:1000 dilution. Alexa fluor secondary antibodies (A10521, A10520, A-21236, and A-21245) were all purchased from Thermo Scientific and used at 1:1000 dilutions. IL6 FISH mRNA probe set (VA6-12712-VC) was purchased from Thermo Scientific. CF 488A succinimidyl ester (SCJ4600018) was purchased from Sigma and used at a 1:10,000 dilution. Hoechst 33342 was purchased from Thermo Scientific (H3570) and used at a 1:10,1000 dilution. The Senescence β-Gal Staining Kit was purchased from CST (9860). The ViewRNA ISH Cell Assay Kit was purchased from Thermo Scientific (QVC0001). Etoposide (E1383) and chloroquine were purchased from Sigma (AAJ6445914). Palbociclib (S1116) and trametinib (S2673) were purchased from Selleckchem.

### Cell lines and culture media

MCF10A (ATCC CRL-10317) cells were obtained from ATCC and grown in DMEM/F12 supplemented with 5% horse serum, 20 ng/ml EGF, 10 μg/ml insulin, 0.5 μg/ml hydrocortisone, 100 ng/ml cholera toxin, and 100 μg/mL of penicillin and streptomycin. MCF10A starvation media consisted of DMEM/F12, 0.5 μg/ml hydrocortisone, 100 ng/ml cholera toxin, and 100 μg/mL of penicillin and streptomycin. During live-cell imaging, phenol red-free full growth media was used. RPE-hTERT (ATCC CRL-4000) were obtained from ATCC grown in DMEM/F12 supplemented with 10% FBS, 1x Glutamax, and 100 μg/mL of penicillin and streptomycin. MCf7 (ATCC HTB-22) were obtained from ATCC and grown in RPMI supplemented with 10% FBS, 1x Glutamax, and 100 μg/mL of penicillin and streptomycin. WI38-hTERT cells were grown in DMEM supplemented with 10% FBS and 100 μg/mL of penicillin and streptomycin. All cell lines were grown in a humidified incubator at 5% CO2 and 37 °C.

### Drug treatments

MCF10A cells were plated at 100,000 cells per well in a plastic 6 well culture plate before being treated with 10 μM etoposide the following day for 24h. Etoposide was removed and cells were washed once with PBS before being returned to full growth media. The cells were maintained in culture throughout the duration of drug recovery with media refreshes every 3d. 24h prior to imaging, the etoposide-released cells were trypsinized and replated onto a collagen coated (1:50 dilution in water) (Advanced BioMatrix, No. 5005) 96-well glass-bottom plate (Cellvis Cat. No. P96-1.5H-N) at 1500 cells per well for live-cell imaging and 3000 cells per well for immunofluorescence. To induce quiescence, 1500 cells per well were plated directly onto a collagen coated 96-well glass-bottom plate and treated continuously with 3 μM Palbociclib, 100 nM Trametinib, or serum free media for up to 12 days. Contact inhibited cells were plated at 10,000 cells per well in full-growth media and cultured for up to 12 days. Media was refreshed on all the conditions every 3 days. To perturb autophagy, MCF10A cells were treated with 50 μM chloroquine 3h prior to fixing and staining.

### Flow cytometry

MCF10A cells endogenously tagged with mCitrine-Ki67 and expressing H2B-mTurquoise and DHB-mCherry were trypsinized and resuspended in PBS + 1% FBS + 100 μg/mL of penicillin and streptomycin after a 5d recovery from a 24h treatment with 10 μM etoposide or 10 Gy of ionizing radiation. Unlabeled wild-type cells were used to gate Ki67^off^ cells, which resulted in 25% of etoposide treated cells and 10% of IR treated cells being sorted and replated directly onto a collagen coated (1:50 dilution in water) (Advanced BioMatrix, No. 5005) 96-well glass-bottom plate (Cellvis Cat. No. P96-1.5H-N) for live-cell imaging that started the following day. As a control, the bottom 7.7% of untreated cells were also sorted and plated. For measuring SA-β-Gal in spontaneously quiescent cells, the bottom 1% of mCitrine-Ki67 was sorted and replated as described above for live-cell imaging that started 48h later.

### Immunofluorescence

MCF10A cells were treated with 10 μM etoposide for 24h, washed, and allowed to recover before being seeded onto a collagen coated (1:50 dilution in water) (Advanced BioMatrix, No. 5005) 96-well glass-bottom plate (Cellvis Cat. No. P96-1.5H-N) 24h prior to fixation for 15 minutes with 4% PFA in PBS. Cells were permeabilized at room temperature with 0.1% TritonX for 15 minutes and blocked with 3% Bovine Serum Albumin (BSA) for 1h. Primary antibodies were incubated overnight in 3% BSA at 4 °C and secondary antibodies were incubated for 1-2h in 3% BSA at room temperature. Nuclei were labelled with Hoechst at 1:10,000 in PBS at room temperature for 15 min. Cytoplasms were labelled with succinimidyl ester 488 at 1:10,000 in PBS at room temperature for 30 minutes. Two 100 uL per well PBS washes were performed between each described step. All images were obtained using a 10x 0.4 numerical aperture objective on a Nikon TiE microscope.

### Time-lapse microscopy

MCF10a cells were plated 24h prior to imaging and full growth media was replaced with phenol red-free full-growth media. Images were taken in CFP, YFP, and mCherry every 12 minutes at two sites per well that were spaced 2 mm apart. Total light exposure was kept below 600 ms. Cells were imaged in a humidified, 37°C chamber at 5% CO2. All images were obtained using a 10x 0.4 numerical aperture objective on a Nikon TiE microscope.

### Image processing

Image processing and cell tracking were performed as previously described (Gookin et al., 2017). Phosho-Rb was separated into high and low modes by using the saddle-point in the data as the cutoff (Fig. S1A). Ki67^off^ cells were classified as those less than the 95^th^ percentile of the median nuclear signal in WT cells. Quantification of 53BP1 puncta was determined using a previously described approach (17). Nuclear signals were quantified from a nuclear mask, which was generated using Otsu’s method on cells stained for Hoechst. Cytoplasmic signals were quantified from a cytoplasmic mask, which was generated using Otsu’s method on cells stained for succinimidyl ester. The regionprops function in MATLAB was used to quantify the mean signal for each stain from binary masks of the nucleus or cytoplasm. Immunofluorescence and SA-β-Gal signals were linked back to live-cell imaging traces by nearest neighbor screening after jitter correction as described in (Gookin et al., 2017).

### SA-β-Gal quantification

Compound immunofluorescence + RGB images were obtained by mounting a LIDA light engine attachment to our Nikon TiE widefield microscope and exporting all stacked image channels from ND2 to TIFF via Nikon Elements Viewer. The SA-β-Gal stain for each cell is quantified by measuring the 5^th^ percentile of the cytoplasmic red pixel intensity from pseudo-RGB images of the colorimetric stain (**Fig. 2A**). Cytoplasmic pixels were indexed from the binary mask generated with succinimidyl ester 488 as described above.

SA-β-Gal staining intensity is sensitive to the cell fixation method; 2% PFA and SA-β-Gal CST kit fixatives were compared (**Fig. S2A-C**). Although the dynamic range of SA-β-Gal staining is larger for the kit fixative compared to 2% PFA, the kit fixative is less compatible with subsequent immunofluorescence staining (excluding LAMP1) (**Fig. S2C**). The kit fixative was used for SA-β-Gal staining following all live-cell imaging experiments and LAMP1 immunofluorescence. 2% PFA was used for all other SA-β-Gal + immunofluorescence experiments.

To validate the SA-β-Gal quantification method, the upper and lower quartiles of SA-β-Gal intensities were displayed through a binary cytoplasmic mask filter that was gated from the distribution of SA-β-Gal values after a 4d release from a 24h pulse 10 μM etoposide (**Fig. S3A**).

Immunofluorescence co-staining with SA-β-Gal is limited to the Cy5 channel due to strong bleed-through fluorescence in the GFP channel and partial bleed-through into the Cy3 channel after staining with SA-β-Gal (kit fixative) (**Fig. S3B**). Cy3 was only used for phospho-Rb (S807/811) co-staining since the bimodality of the phospho-Rb distribution is well maintained even after SA-β-Gal staining in the 2% PFA condition (**Fig. S2C**).

### Receiver operating characteristic (ROC) analysis

We performed a ROC analysis by determining the false positive and true positive rate of detection for SA-β-Gal, LAMP1, and cytoplasmic area by sliding the cutoff at every 10^th^ percentile of intensity. The cutoff for 53BP1 was at increasing numbers of nuclear bodies (from 0 to 8+ foci). Cells were classified as proliferating, quiescent, or senescent. Our ground-truth definition for senescence required that cells remained CDK2^low^ throughout the duration of 4 days of live-cell imaging and was used to calculate false and true positive rates.

### Statistical analyses

All statistical tests shown are two-sample *t*-tests: *p < 0.05; **p < 0.01, ***p < 0.001, ****p < 0.0001 (**Table S1**). All error bars represent the standard error of the mean derived from multiple technical replicates (**Table S2**). Technical replicates shown for all experiments come from data representative of at least two biological replicates.

## Data availability

All data will be deposited to a publicly available repository. No original code is reported in this paper.

## Acknowledgments

We thank Judith Campisi for providing us with WI38-hTERT immortalized cells. We thank Theresa Nahreini for her expertise and assistance with cell sorting. We thank Joe Dragavon for his insights on quantitative microscopy. We thank Steve Cappell and Iain Cheeseman for reading the manuscript pre-submission. This work was supported by an NIH Training Grant T32 (5T32GM008759-19 (to H.M.A)) and an NIH Director’s New Innovator Award (Award 1DP2CA238330-01 (to S.L.S)).

## Author contributions

H.M.A. designed research; H.M.A. and B.F. conducted research; H.M.A. analyzed data; H.M.A. and S.L.S conceived the project; and H.M.A. and S.L.S wrote the paper; S.L.S supervised the project.

## Declaration of interests

The authors declare no competing interests.

---

**Supplemental Figure 1.**
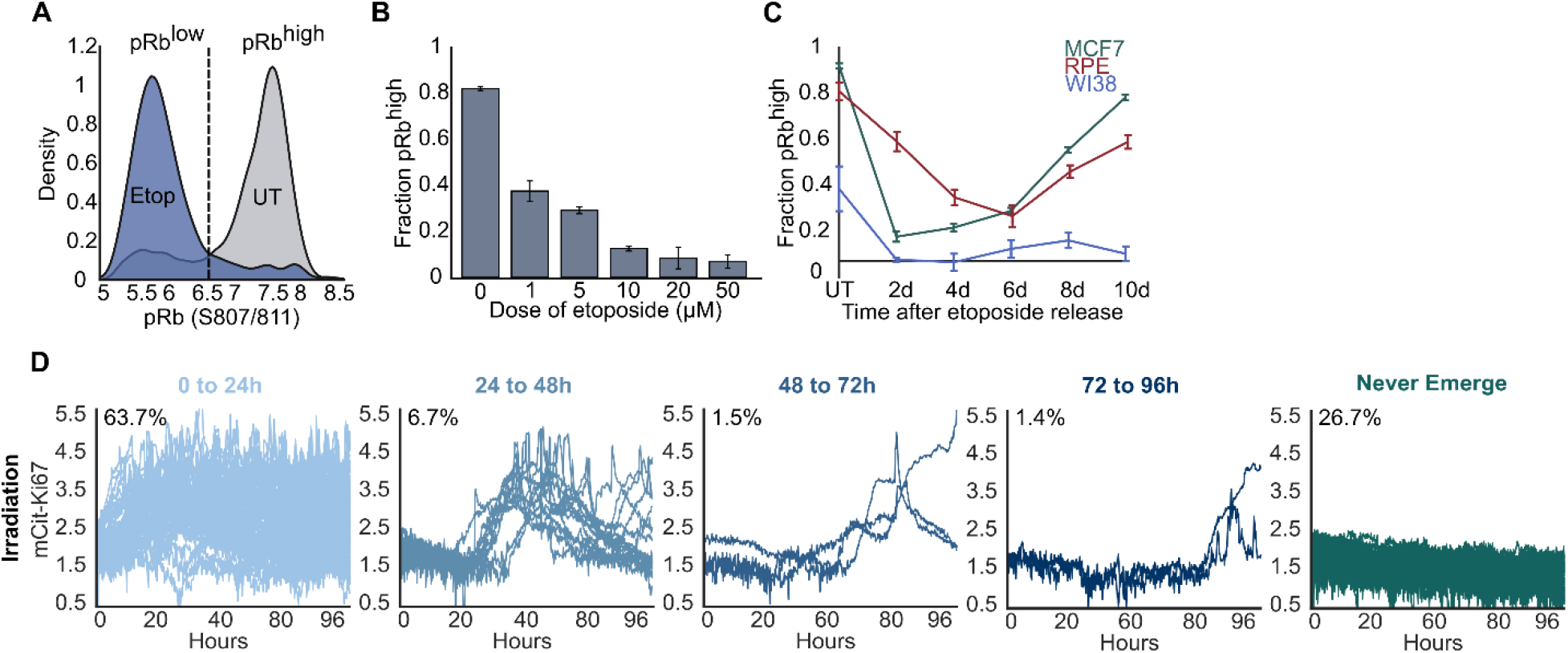
Release from acute chemotherapeutic stress universally induces population heterogeneity in cell cycle fate. (A) Definition of phospho-Rb^low^ versus phospho-Rb^high^ in MCF10A cells released from a 24h treatment of 10 μM etoposide for 6d. (B) MCF10A cells were treated with increasing concentrations of etoposide for 24h before being fixed and stained for phospho-Rb at 6d after release from drug. (C) MCF7, RPE, and WI38-hTERT cells were treated with 10 μM etoposide for 24h before being washed and allowed to recover for up to 10d. (D) MCF10A cells expressing endogenously tagged with mCitrine-Ki67 were plated on day 0 and treated with 10 Gy ionizing radiation the following day. On day 5, Ki67^off^ cells were isolated by flow cytometry, plated, and allowed to grow for 24h before being imaged for 96h by time-lapse microscopy. Single-cell traces are grouped based on their relative timing of cell-cycle re-entry from the Ki67^off^ state and the percent of cells in each group is indicated. 200 traces are plotted in total.

**Supplemental Figure 2.**
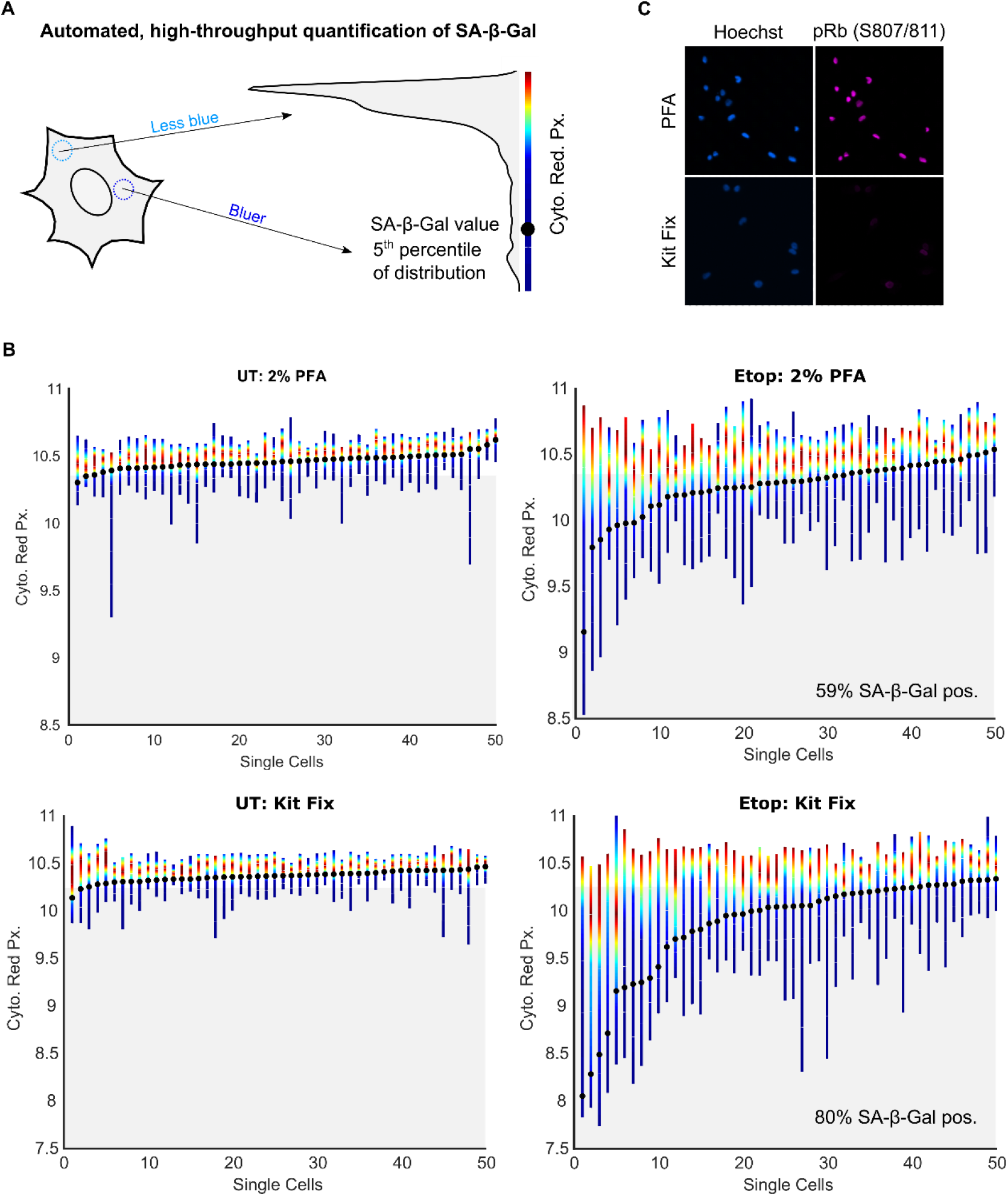
Automated, high-throughput quantification of the SA-β-Gal stain in single cells induced to senescence. (A) A single representative cell’s cytoplasmic red pixel distribution. The heatmap corresponds to the relative frequency of events along the distribution. (B) Quantification of SA-β-Gal for 50 single cells left untreated or released for 6d from a 24h treatment with 10 μM etoposide, for both 2% PFA (which is optimal for immunofluorescence + SA-β-Gal) and the CST kit fixative (which is optimal for detection of SA-β-Gal). The black dot is the value at the 5^th^ percentile of the distribution, which is the SA-β-Gal score for that cell. The percentage of SA-β-Gal^pos^ cells was calculated using the 95^th^ percentile of all untreated cells as the cutoff. Heatmap coloring as in panel A. (C) Comparison of immunofluorescence for phospho-Rb (S807/811) after fixation with 2% PFA versus CST SA-β-Gal kit fixative in WT MCF10A cells showing the weaker immunofluorescence signal with the CST kit fixative.

**Supplemental Figure 3.**
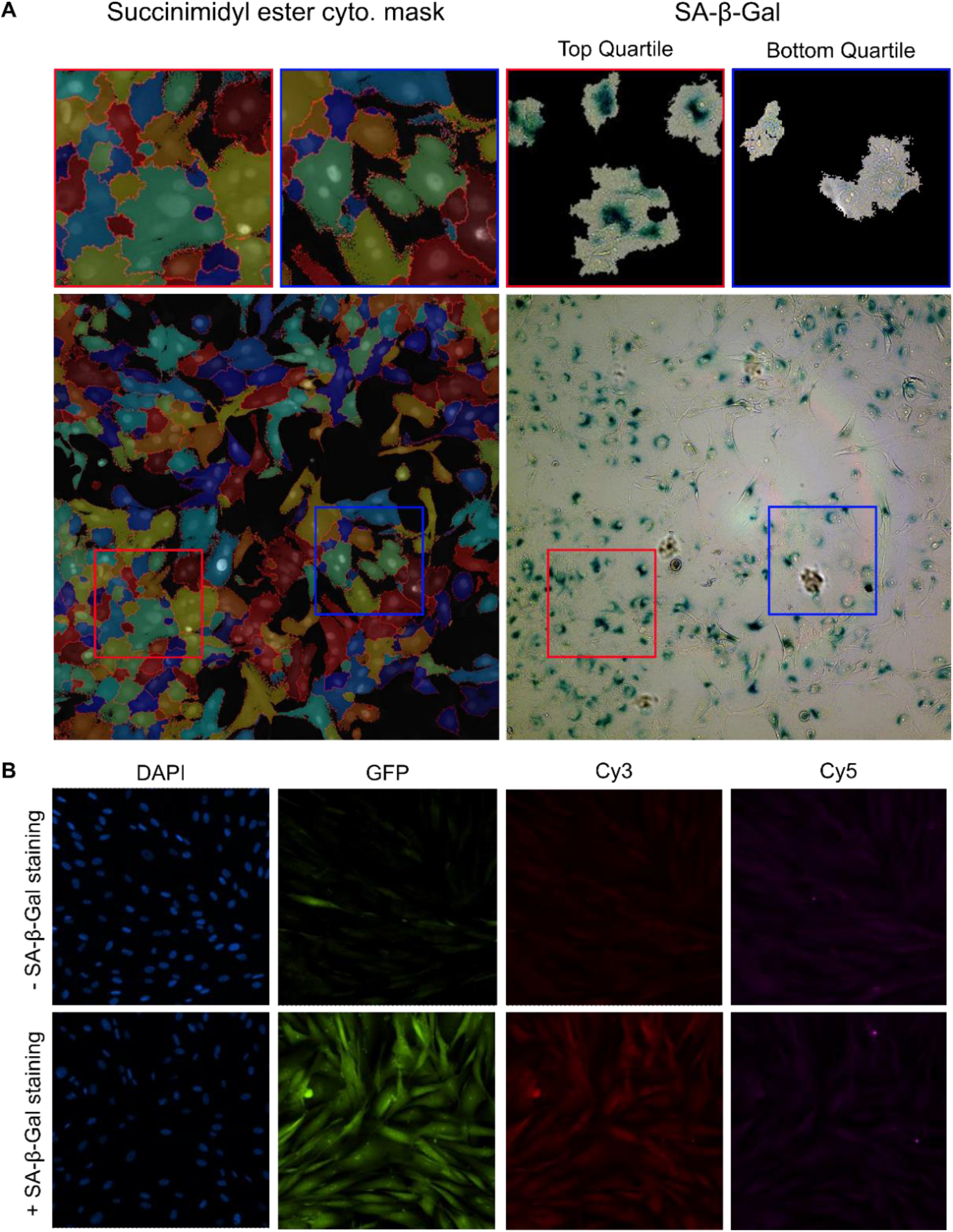
Validation of the SA-β-Gal quantification method and comparison of background fluorescence. (A) Validation of whole-cell segmentation using the succinimidyl ester stain and SA-β-Gal quantification by displaying the upper and lower quartiles of SA-β-Gal signals from the binary cytoplasmic mask after a 4d release from a 24h pulse of 10 μM etoposide in MCF10A cells. (B) Comparison of background fluorescence after secondary antibody staining in each fluorescent channel with and without co-staining MCF10A cells for SA-β-Gal (kit fixative).

**Supplemental Figure 4.**
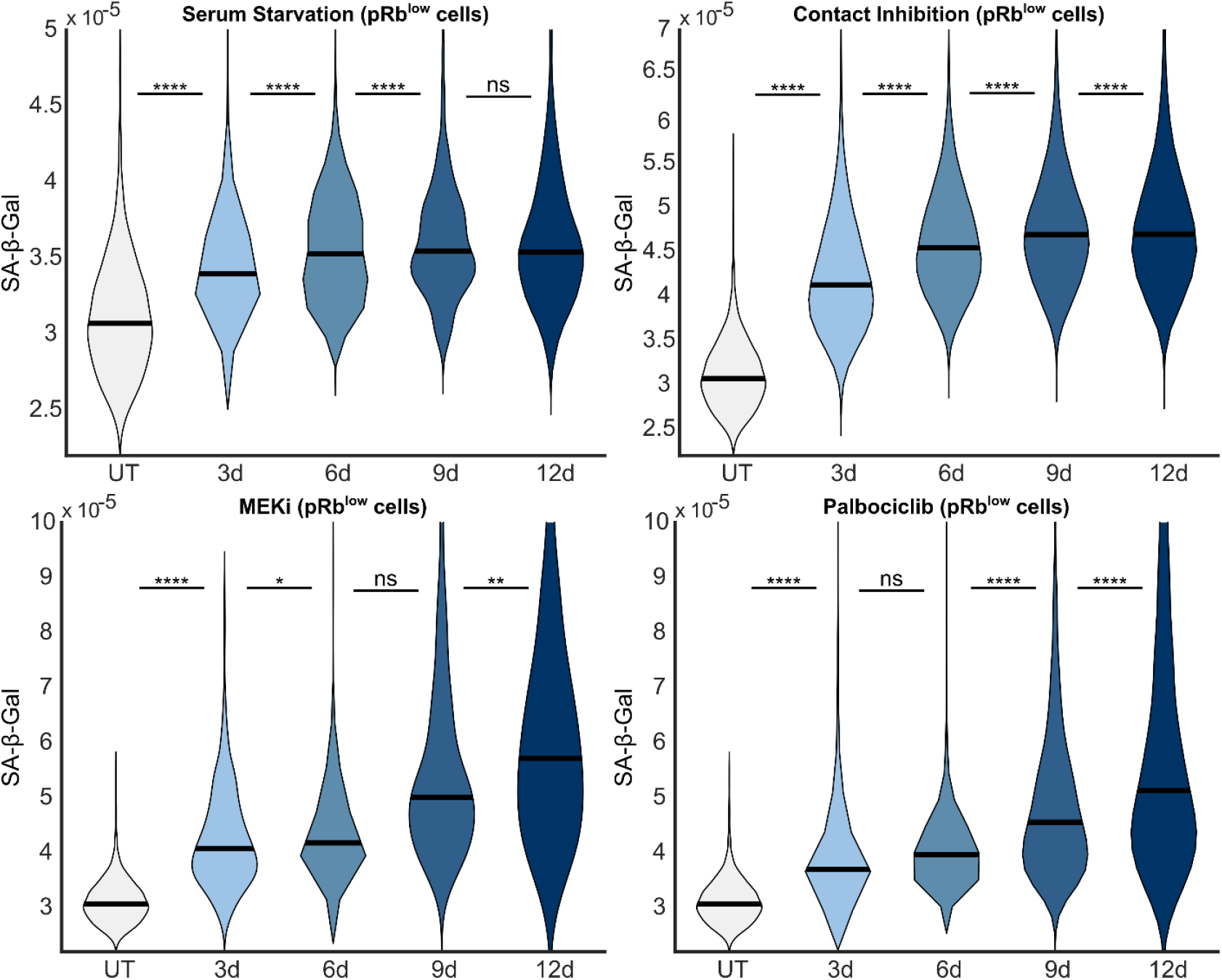
SA-β-Gal staining in phospho-Rb^low^ cells after increasing durations of quiescence induction. Violin plots of the data plotted in Fig. 3E.

**Supplemental Figure 5.**
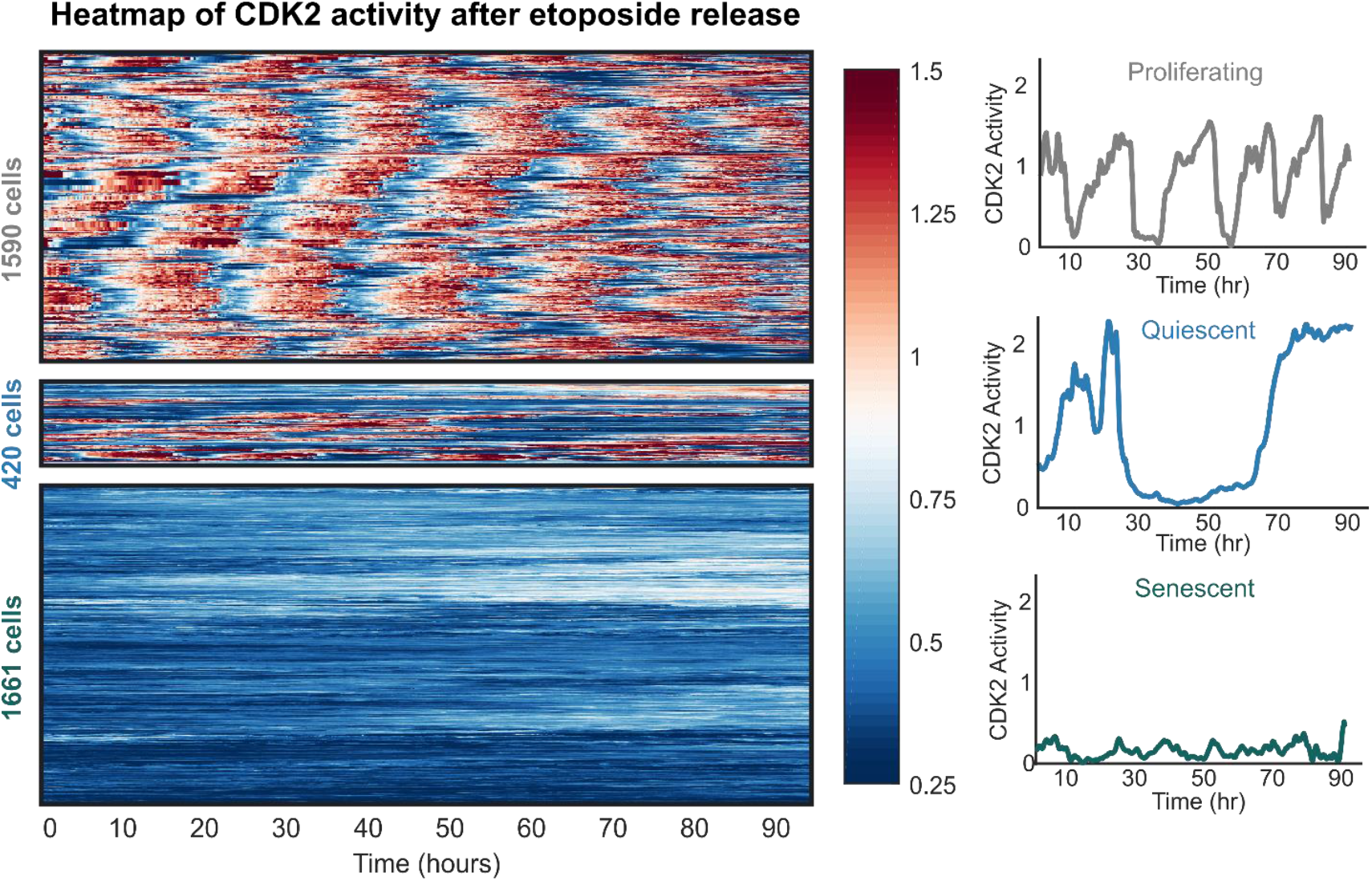
Classification of proliferating, quiescent, and senescent cells. Heatmap of data plotted in Fig. 4A.

## Supplemental Tables

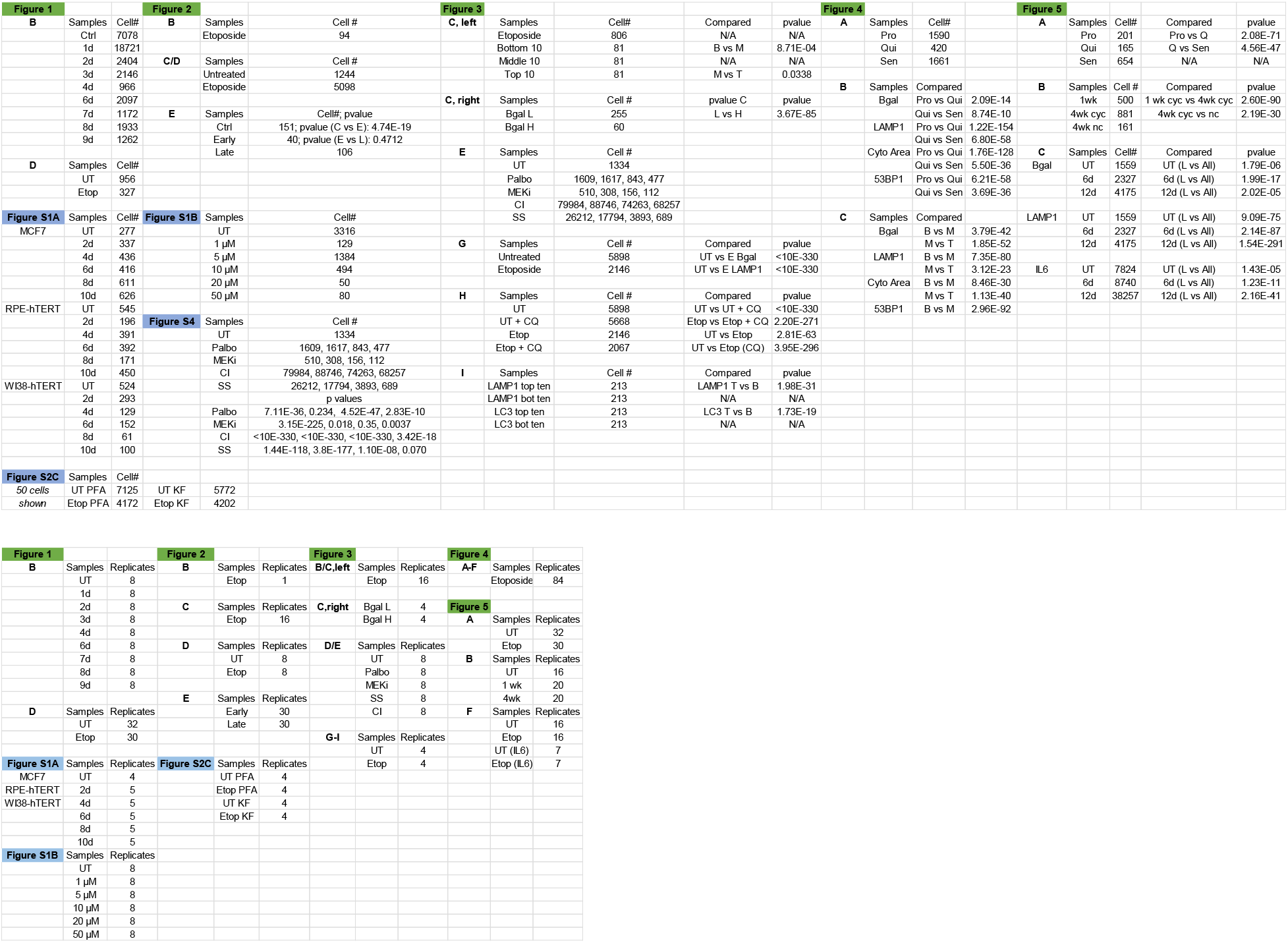

## Notes

### Competing Interest Statement

The authors have declared no competing interest.

